# Strain level centromere variation influences CENP-A association dynamics and centromere function

**DOI:** 10.1101/2022.05.17.492352

**Authors:** Uma P. Arora, Beth A. Sullivan, Beth L. Dumont

## Abstract

Centromeres are rapidly evolving chromatin domains that fulfill essential roles in chromosome segregation. Rapid centromere sequence evolution imposes strong selection for compensatory changes in centromere-associated kinetochore proteins, leading to striking co-evolutionary trends across species. However, it remains unknown whether within species centromere sequence diversity leads to functional differences in kinetochore protein association. House mice (*Mus musculus*) exhibit significant variation in centromere satellite array size and sequence heterogeneity, but the amino acid sequence of CENP-A, a centromere-specific histone variant that specifies centromere identity, is conserved. We hypothesize that centromere satellite sequence variation leads to differences in the localization of CENP-A among house mice, with potential consequences for meiotic drive and genome stability. Using CENP-A chromatin immunoprecipitation with a customized *k*-mer based, reference-blind bioinformatic analysis strategy, we compare the CENP-A sequence association landscape in four diverse inbred mouse strains (C57BL/6J, CAST/EiJ, LEWES/EiJ, and PWK/PhJ). We uncover significant strain-level diversity in CENP-A associated sequences, with more closely related strains exhibiting more similar CENP-A association profiles. LEWES/EiJ and CAST/EiJ show mild association of CENP-A with the pericentromeric satellite repeat, countering the prevailing notion that functional centromere size is solely determined by the size of the minor satellite array. Strain-specific CENP-A association profiles are enriched for unique suites of transcription factor motifs, hinting at strain differences in centromere transcription. Given the importance of centromere-CENP-A association and centromere transcription for both kinetochore assembly and chromosome segregation fidelity, our findings suggest a potential mechanism for centromere-mediated variation in genome stability among inbred mouse strains.

## INTRODUCTION

Centromeres are satellite-rich chromatin domains that serve as sites for the assembly of the multiprotein kinetochore complex which facilitates chromosome segregation during mitosis and meiosis (Bakhoum et al. 2009; Fukagawa and Earnshaw 2014; McKinley and Cheeseman 2016; Schalch and Steiner 2017). Loss of centromere integrity and function can lead to apoptosis, chromosome mis-segregation, and widespread genome instability, phenomenon intimately associated with cancer and infertility (Giunta and Funabiki 2017; Shrestha et al. 2021).

Despite their essential roles, centromeres are highly variable in sequence and structure between and within species (Musich et al. 1980; Alexandrov et al. 2001; Ventura et al. 2007; Malik and Henikoff 2009; Alkan et al. 2011; Rocchi et al. 2012; Cacheux et al. 2016; Arora et al. 2021). Rapid sequence level evolution of centromeres imposes strong evolutionary pressures for cognate changes to DNA-associated kinetochore proteins. As a result, several kinetochore proteins are adaptively co-evolving with centromere satellite DNA (Malik and Henikoff 2001; Talbert et al. 2004). The most well-established case involves CENP-A, a H3 histone variant that delimits the core centromere domain, helps maintain centromere repeat stability, and directs assembly of the kinetochore complex (Foltz et al. 2006; Okada et al. 2006; Perpelescu and Fukagawa 2011; Fukagawa and Earnshaw 2014; Giunta and Funabiki 2017). Although its homolog, histone H3, has been subject to intense purifying selection over evolutionary time and is highly conserved across taxa, CENP-A exhibits clear signals of adaptive evolution within its DNA-associating NH_2_-terminal tail (Henikoff et al. 2001; Malik and Henikoff 2001; Talbert et al. 2004; Malik 2009; Malik and Henikoff 2009; Drinnenberg et al. 2016). The molecular arms race between CENP-A and centromere satellites preserves the compatibility of this protein-DNA association, safeguarding assembly and function of the kinetochore.

Centromeres persist as gaps in most high-quality mammalian reference genomes. The absence of high-quality sequence information poses a barrier to the characterization of centromere variation and its potential functional consequences for kinetochore assembly and chromosome segregation. Recently, we used *k*-mer based bioinformatic methods in conjunction with cytogenetic approaches to uncover remarkable variation in the size and sequence composition of centromeres across diverse inbred mouse strains and wild-caught house mice (*Mus musculus*) (Arora et al. 2021). In house mice, each centromere can be broadly categorized into two chromatin domains. The core functional centromere domain is comprised of a single 120-bp minor satellite repeat unit that is iterated in tandem over ∼1Mb of sequence. This minor satellite array facilitates CENP-A association and kinetochore assembly at the centromere. The minor satellite array is flanked by a ∼2Mb pericentromeric chromatin domain defined by a focal 234-bp major satellite repeat monomer and the heterochromatin mark H3K9me3. Our previous work showed that inbred mouse strains differ in the sequence composition of their resident minor satellite repeats and exhibit variable levels of minor satellite sequence heterogeneity. The potential functional consequences of this sequence diversity for the dynamics of centromeric chromatin and kinetochore assembly remain to be elucidated. Importantly, even subtle perturbations to kinetochore assembly or stability could have downstream impacts on the fidelity of chromosome segregation and propensity for centromere drive (Henikoff et al. 2001; Malik and Henikoff 2001; Pardo-Manuel de Villena and Sapienza 2001; Talbert et al. 2004; Kursel and Malik 2018; Kruger and Mueller 2021; Kumon et al. 2021).

Despite significant sequence variation in centromere repeats across house mice, there are no known amino acid differences in CENP-A within *Mus musculus*. This conservation raises the question of whether CENP-A exhibits sequence-specific binding in *Mus musculus* strains with divergent centromere satellite DNA. Prior work suggests that CENP-A preferentially associates with centromere satellite repeat arrays with lower repeat diversity (Aldrup-MacDonald et al. 2016). For instance, in maize, different alternating centromere satellite monomers position CENH3 (the plant ortholog of CENP-A) with translational and rotational phasing to ensure regular spacing of nucleosomes on each monomer (Zhang et al. 2013). Thus, the strain-specific organization of centromere satellite repeat diversity could influence CENP-A spacing and, in turn, modulate kinetochore complex density and impact genome stability (Walstein et al. 2021).

The extensive variation in centromere satellites across inbred mouse strains provides a powerful experimental framework to explicitly test how centromere sequence variation influences CENP-A association (Arora et al. 2021). To this end, we performed CENP-A chromatin immunoprecipitation followed by sequencing (ChIP-seq) across four diverse inbred mouse strains: C57BL/6J (reference strain, *M. m. domesticus*), LEWES/EiJ (*M. m. domesticus*), CAST/EiJ (*M. m. castaneus*), and PWK/PhJ (*M. m. musculus*). These strains capture the full breadth of inbred mouse centromere minor satellite heterogeneity and provide sampling across the three principal *Mus musculus* subspecies. Using both consensus-guided and reference independent computational analyses, we uncover divergent CENP-A sequence association landscapes between strains. Overall, our findings point to strain differences in functional centromere size, CENP-A density, and transcription factor association, hinting at potential strain differences in kinetochore protein assembly and chromosome segregation dynamics.

## RESULTS

### Validation of CENP-A ChIP in diverse inbred strains

Mammalian centromere chromatin is defined by interspersed blocks of histone H3 and CENP-A, the centromere-specific histone H3 variant. We sought to characterize strain differences in CENP-A sequence association across four diverse inbred *Mus musculus* strains: C57BL/6J, LEWES/EiJ, CAST/EiJ, and PWK/PhJ (Figure 1A). To confirm the efficacy of immunoprecipitation in each inbred strain, we quantified the enrichment of mononucleosomal DNA sequences after immunoprecipitation for CENP-A, H3K4me3 (positive control for euchromatic genomic regions), and IgG (negative control) compared to input mononucleosomal DNA using qPCR (Supplemental Figure S1). We then sequenced the input (3 replicates) and chromatin immunoprecipitated DNA with antibodies targeting CENP-A (3 replicates), H3K4me3 (1 replicate), and IgG (1 replicate) for each strain (except C57BL/6J with 2 instead of 3 replicates for CENP-A ChIP and input).

**Figure 1:**
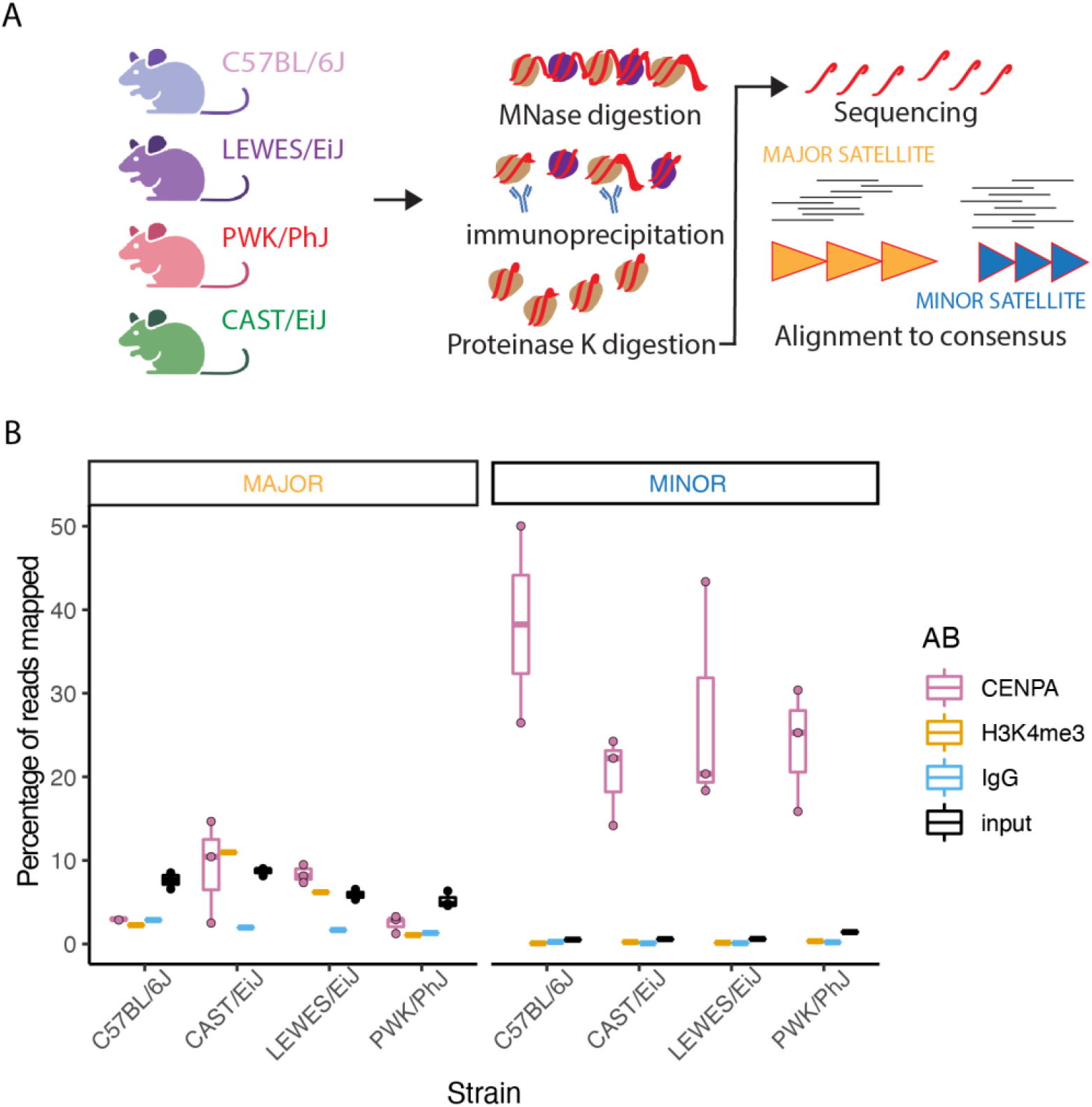
Schematic and validation of CENP-A ChIP-seq across diverse inbred strains. (A) Overview of the experimental design of the ChIP-seq experiments performed and the consensus-guided genomic analyses for CENP-A ChIP sequence enrichment. (B) Boxplots representing the percentage of reads that mapped to either the major satellite (left) or minor satellite (right) consensus sequences. Color represents sample identity (AB: antibody). For each boxplot, the horizontal line represents the mean, the top and bottom of the box represent the upper and lower quartile respectively, and the vertical line represents the range of values.

To assess the efficiency of CENP-A sequence enrichment in each inbred strain, we estimated the percentage of sequencing reads that mapped to the major and minor centromere satellite consensus sequences derived from the C57BL/6J inbred mouse strain (Figure 1B). Prior work has showed that CENP-A primarily associates with the minor satellite, with negligible CENP-A association with the pericentromere-enriched major satellite (McKinley and Cheeseman 2016). As expected, a higher proportion of sequencing reads map to the minor satellite consensus sequence, as opposed to the major satellite consensus sequence, in CENP-A ChIP compared to input samples (Wilcox rank sum exact test *P* = 2.83×10^-6^; Figure 1B). In addition, the percentage of reads that mapped to the minor satellite consensus was not significantly different across strains indicating a uniform enrichment of minor satellite consensus sequences from CENP-A ChIP across strains (Kruskal Wallis one way ANOVA *P* = 0.95; Figure 1B, right). As expected, fewer reads from the IgG negative control ChIP map to either the major or minor satellite consensus sequences. Taken together, these results confirm that CENP-A ChIP enriches for centromere-associated minor satellite sequences across all four inbred strains.

### Variable CENP-A-associated consensus sequence enrichment across strains

To investigate strain differences in CENP-A/DNA association, we identified CENP-A associated sequences with the most variable enrichment across strains (Supplemental Figure S2). The canonical 120 bp minor satellite consensus repeat and a shorter version with an 8 bp deletion between positions 82-89 were the most differentially abundant sequences between strains (Supplemental Figure S2). These two sequences comprise the two most dominant minor satellite variants in C57BL/6J, the reference strain from which the consensus sequence was originally derived (Rice 2020).

Compared to the other three strains, C57BL/6J exhibits the greatest relative enrichment of CENP-A ChIP sequenced reads mapping to the reference minor satellite consensus (ANOVA *P* < 0.01). C57BL/6J is an inbred laboratory strain of predominantly *M. m. domesticus* origin. LEWES/EiJ, an inbred strain developed from wild-caught *M. m. domesticus* in Delaware, USA, showed the second highest level of enrichment. The two strains of non-domesticus subspecies origin (CAST/EiJ (*M. m. castaneus*) and PWK/PhJ(*M. m. musculus*)) have reduced levels of CENP-A association with the reference minor satellite sequence. We previously showed that the repertoire of minor satellite sequences varies across diverse mouse strain genomes (Arora et al. 2021). Our current findings extend this result to show that CENP-A differentially associates with specific minor satellite sequences in these strains.

### Strain variation in the relative enrichment of CENP-A at pericentromeric chromatin

We observe differential enrichment of CENP-A ChIP reads mapping to the major satellite consensus sequence across strains (Figure 1B). In strains C57BL/6J and PWK/PhJ, fewer reads map to the major satellite in CENP-A ChIP compared to input samples (Figure 1B, left). This pattern accords with expectations, as CENP-A is thought to primarily associate with the minor satellite (McKinley and Cheeseman 2016). However, strains CAST/EiJ and LEWES/EiJ show an opposite pattern, with an enrichment of reads mapping to the major satellite relative to input samples (Kruskal Wallis one-way ANOVA *P* = 0.07; Figure 1B, left). These intriguing findings suggest that CENP-A may associate with the major satellite in some inbred mouse strains. Thus, minor satellite array size might not be the only genetic factor influencing functional centromere size, as previously reported in mice (Iwata-Otsubo et al. 2017).

We performed one replicate H3K4me3 immunoprecipitation per strain as a positive control for euchromatic, non-centromeric regions. Previous research suggests that H3K4me3 is not found in the core centromere region (minor satellite) (Sullivan and Karpen 2004; Li et al. 2008), but there is minimal information regarding its enrichment in pericentromeric chromatin. In agreement with data from the literature, we find no evidence of H3K4me3 ChIP reads mapping to the minor satellite consensus sequence in any strains (Figure 1B, right). However, strains CAST/EiJ and LEWES/EiJ exhibited greater H3K4me3 ChIP signal across the major satellite than C57BL/6J and PWK/PhJ, a strain trend that follows the pattern observed with CENP-A enrichment (Figure 1B, left). Our findings raise the intriguing possibility of strain differences in specific epigenetic modifications of the major satellite domain that could influence the chromatin landscape and CENP-A localization at the centromere, although more replicates are required to make any conclusions.

### Strain differences in CENP-A positioning along the minor satellite repeat

To investigate strain variation in CENP-A positioning along the minor satellite consensus, we profiled relative read mapping coverage along the minor satellite consensus sequence in CENP-A ChIP compared to input samples. Each strain exhibits a unique spatial pattern of CENP-A positioning (Figure 2A; Kruskall-Wallis; df =3, *P* =0.0019). C57BL/6J and PWK/PhJ share a common peak between positions 65-75. These sites overlap the CENP-B box, a 17-bp motif that confers sequence-specific binding of the kinetochore protein CENP-B (Masumoto et al. 1989; Ohzeki et al. 2002). CENP-B is a non-essential kinetochore protein (Hudson et al. 1998), but is thought to enhance stability of CENP-A-DNA association and play important roles in de-novo centromere formation (Ohzeki et al. 2002; Fachinetti et al. 2015). We asked whether those strains with CENP-A positioning at the CENP-B box (C57BL/6J and PWK/PhJ) simply had more occurrences of the CENP-B box in their CENP-A ChIP-seq reads. We confirmed that the peak of CENP-A association at the CENP-B box in C57BL/6J and PWK/PhJ is not due to an overall increase in the abundance of CENP-B boxes in CENP-A ChIP-seq reads; similar numbers of CENP-B box motifs were recovered for PWK/PhJ and both LEWES/EiJ and CAST/EiJ, two strains with no peak of CENP-A association at this locus (Figure 2B; Dunn test pairwise comparison *P* > 0.05). Previous work investigating CENP-A positioning along the minor satellite in C57BL/6J and ZALENDE/EiJ, a wild-derived inbred strain of *M. m. domesticus* ancestry harboring several Robertsonian chromosomal fusions, found that CENP-A was centered at the CENP-B box in both strains. Our findings indicate that the CENP-B box does not universally centralize positioning of CENP-A in all inbred mouse strains. Indeed, the most prominent peak for CAST/EiJ in the CENP-A binding landscape is between positions 83-93, immediately adjacent to the CENP-B box.

**Figure 2:**
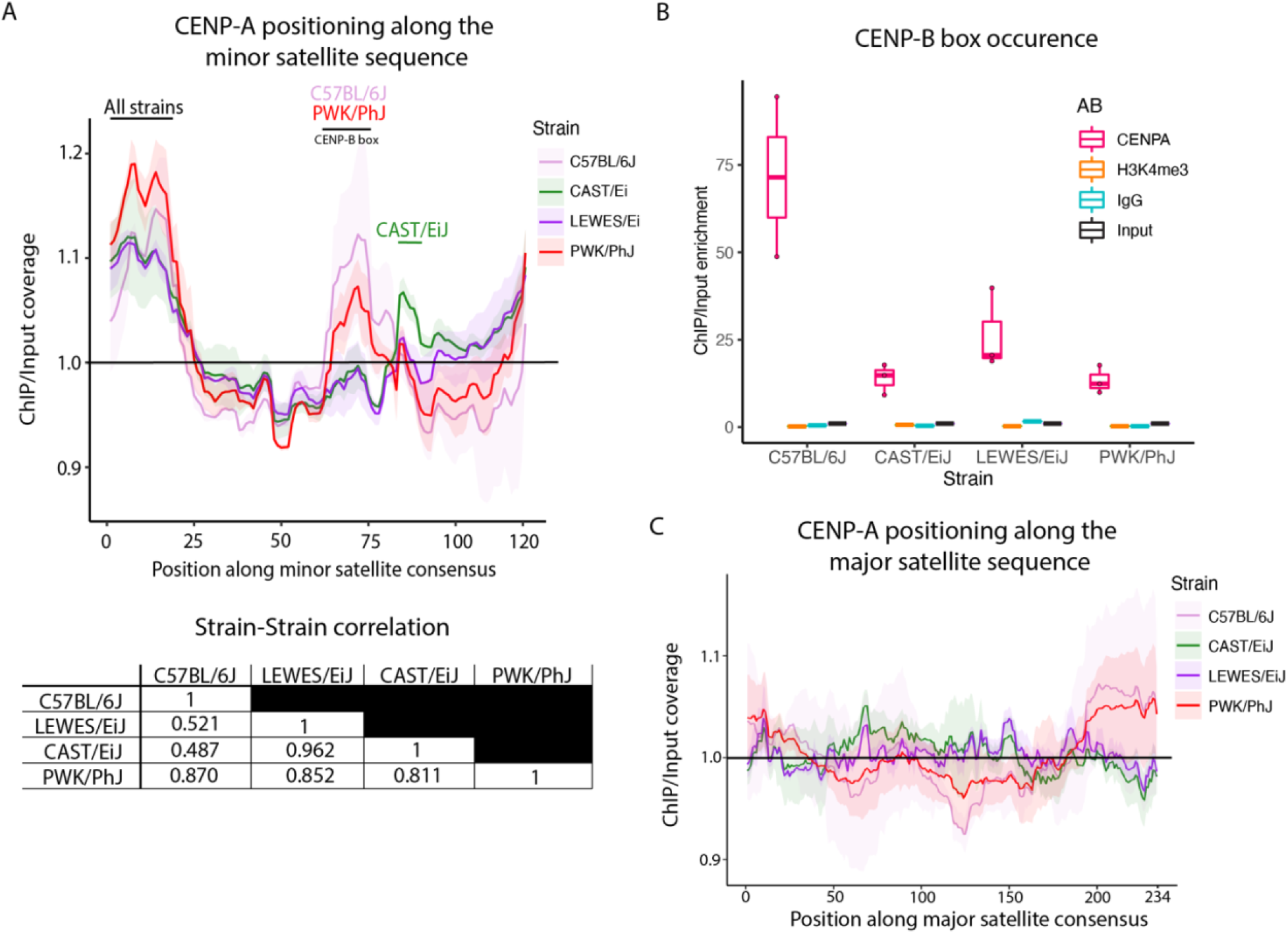
Strain differences in CENP-A positioning and CENP-B motif enrichment. (A) Line plot representing the CENP-A ChIP/input enrichment of read coverage (y-axis) at each position along the minor satellite consensus sequence (x-axis). The shaded region along the line represents +/- 1 standard deviation. The table presents paired strain Pearson correlations for the average CENP-A ChIP/input enrichment pattern along the minor satellite consensus sequence. (B) Boxplots representing the percent enrichment of CENP-B box motif frequency in ChIP/Input samples. For each boxplot, the horizontal line represents the mean, the top and bottom of the box represent the upper and lower quartile respectively, and the vertical line represents the range of values. (C) Line plot representing the CENP-A ChIP/input enrichment of reads (y-axis) mapped at each position along the major satellite consensus sequence (x-axis). The shaded region along the line represents +/- 1 standard deviation.

Pairwise strain correlations among minor satellite CENP-A enrichment profiles indicate that strain pair CAST/EiJ and LEWES/EiJ and strain pair C57BL/6J and PWK/PhJ have more similar CENP-A positioning patterns (Figure 2A; C57BL/6J-PWK/PhJ correlation r^2^ = 0.870, *P* < 2.2×10^-16^; CAST/EiJ-LEWES/EiJ correlation r^2^ = 0.962, *P* < 2.2×10^-16^). This trend counters expectations based on overall strain relatedness. LEWES/EiJ and C57BL/6J share a common principal subspecies designation (*M. m. domesticus*) and a priori might be expected to exhibit a more highly conserved CENP-A association landscape than inter-subspecies comparisons. The absence of such a trend implies the potential for rapid positional changes in CENP-A binding at centromeres. Taken together, our findings suggest that CENP-A associates with distinct sequence-specific contexts within the minor centromere satellite unit in different mouse strains.

CENP-A positioning along the major satellite consensus sequence shows higher strain similarity than observations with the minor satellite (df=3, *P* =0.02214; Figure 2C). However, we acknowledge that with a limited number of reads mapping to the major satellite, we may be underpowered to detect strain differences.

Interestingly, the two strains that demonstrate slight enrichment of major satellite sequences in CENP-A ChIP compared to input (CAST/EiJ and LEWES/EiJ; Figure 1B) also harbor peaks at positions 60-70 and 145-155 along the major satellite consensus sequence (Figure 2C).

### Most CENP-A enriched *k*-mers are strain-specific

The consensus-driven read mapping strategy employed above is potentially biased by its reliance on a reference sequence derived from the C57BL/6J inbred strain. To circumvent this limitation, we adapted a *k*-mer based approach to investigate strain differences in CENP-A associated sequences in a manner agnostic to the reference consensus centromere satellite sequence (Smith et al. 2021). First, we counted the frequency of all *31*-mers observed in each replicate CENP-A ChIP and input sample. Next, for each replicate, we quantified the enrichment of each unique *31*-mer as the ratio of read count normalized 31-mer frequency in CENP-A ChIP relative to input (“enrichment score”; Supplemental Figure S3). Lastly, for each *31*-mer, we averaged the CENP-A ChIP/input enrichment score across replicates and extracted the 0.1% most enriched *31*-mers for each strain. We picked 0.1% as the cutoff to ensure selection of an optimal number of *k*-mers that capture strain-specific sequence features. This approach enables unbiased selection of the most CENP-A enriched *31*-mers in each strain.

We compared these strain-specific sets of CENP-A enriched *31*-mers and observe significantly more *k*-mer sharing between strains than expected by chance (*P* < 1×10^-6^; Figure 3). This finding reveals some underlying conservation of CENP-A associated sequences between diverse strains. However, despite this backdrop of weak conservation, the vast majority of enriched *k*-mers in a strain are unique to each strain (78.8% for C57BL/6J; 76.1% for CAST/EiJ; 82.6% for LEWES/EiJ; 72.4% for PWK/PhJ). Thus, diverse mouse strains harbor libraries of CENP-A associated sequences that are largely non-overlapping with each other.

**Figure 3:**
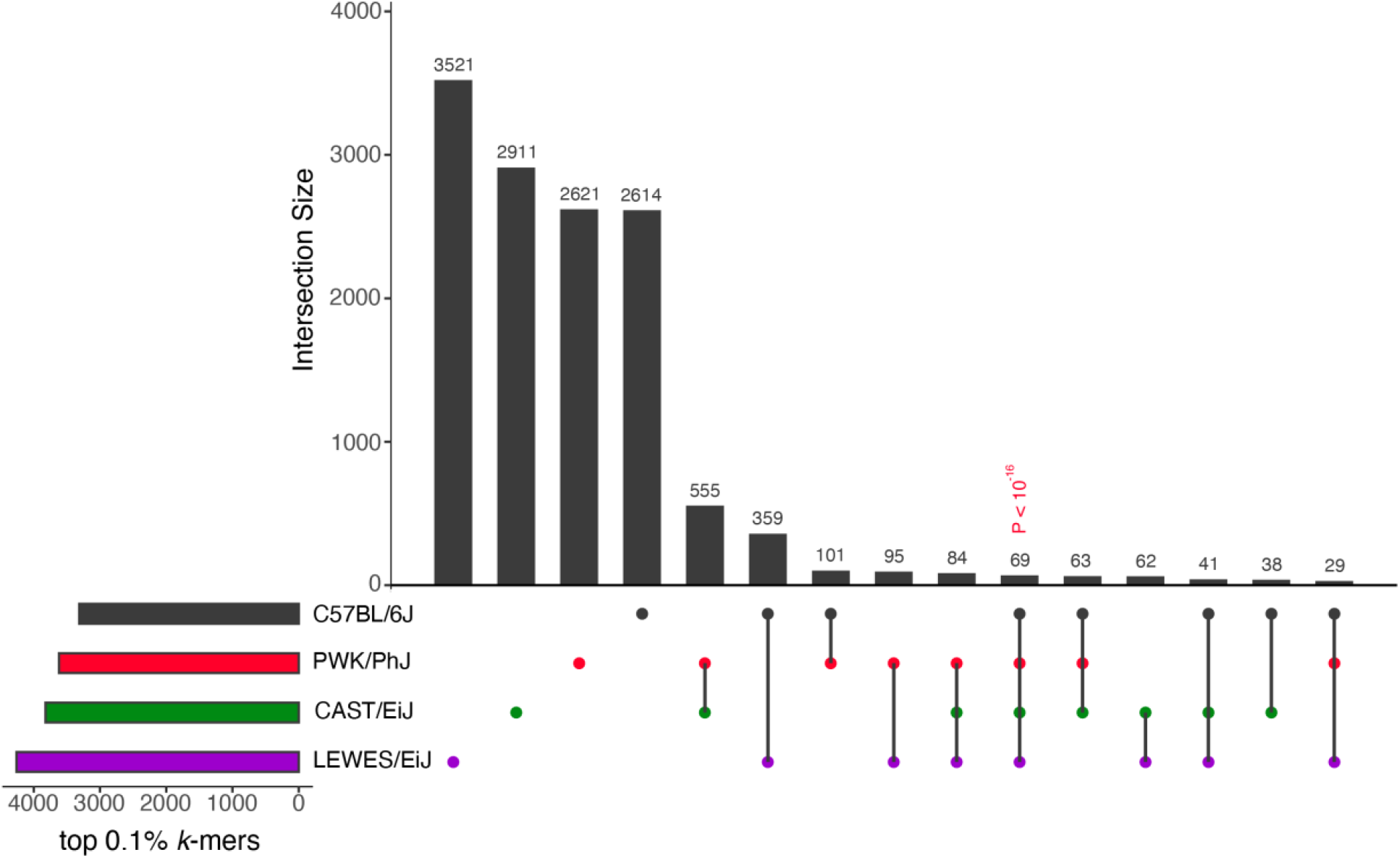
Majority of CENP-A enriched *k*-mers are strain-specific. Upset plot representing the extent of *k*-mer sharing among the top 0.1% most enriched CENP-A/input *k*-mers in each strain. The total number of *k*-mers from each strain’s set of CENP-A enriched *k*-mers is represented by the horizontal bars. The vertical bars represent the number of *k*-mers that belong to one or more strains, indicated by the dots below.

We next compared the set of CENP-A enriched *k*-mers for each strain to the consensus minor satellite sequence. Remarkably, none of these enriched *31*-mers present a perfect match to the minor satellite consensus sequence. Indeed, most enriched *31*-mers harbor two or more sequence mismatches from the consensus minor satellite sequence (Supplemental Figure S4). We conclude that, while the house mouse consensus sequence provides a useful tool to assess relative satellite diversity present across strains, it does not comprehensively represent sequences that preferentially associate with CENP-A in any inbred mouse strain, including C57BL/6J.

### Discovery of strain-specific CENP-A associated sequence landscapes

To analyze strain-specific sets of CENP-A associated sequences, we developed an approach for quantifying the most sequence abundant and *k*-mer enriched CENP-A associated sequences in each strain. Briefly we assigned each CENP-A ChIP-seq read a score based on the number of enriched *k*-mers within the read (top 0.1% enriched *k*-mers; termed read score) and the normalized abundance of the read in the CENP-A ChIP data (termed read count). We then jointly sorted reads by these two criteria and focused on the 1000 top-scoring reads (Supplemental Figure S5). Surprisingly, this approach yielded non-overlapping sequence sets between strains.

We analyzed the relationship among the top 1000 sequences identified in each strain using a Neighbor Joining tree (Figure 4). CENP-A associated sequences cluster into seven broad clades, with multiple sequences from each strain clustering in each clade. In some clades, there is near-equal representation among strains (clades 2, 5, 6), whereas remaining clades (1, 3, 4, 7) exhibit disproportionate representation of sequences from specific strains (Chi-square test, *P* < 0.05). Most clades are distinguished from their sister clades by long branches, implying significant divergence between sequences in distinct clades. Overall, our analyses identify 7 distinct classes of CENP-A associated sequences that are present in all strains, although the functional distinctions between these sequence groups remains to be determined. Additional work is also required to determine whether sequences from specific clades are disproportionately represented on individual chromosomes.

**Figure 4:**
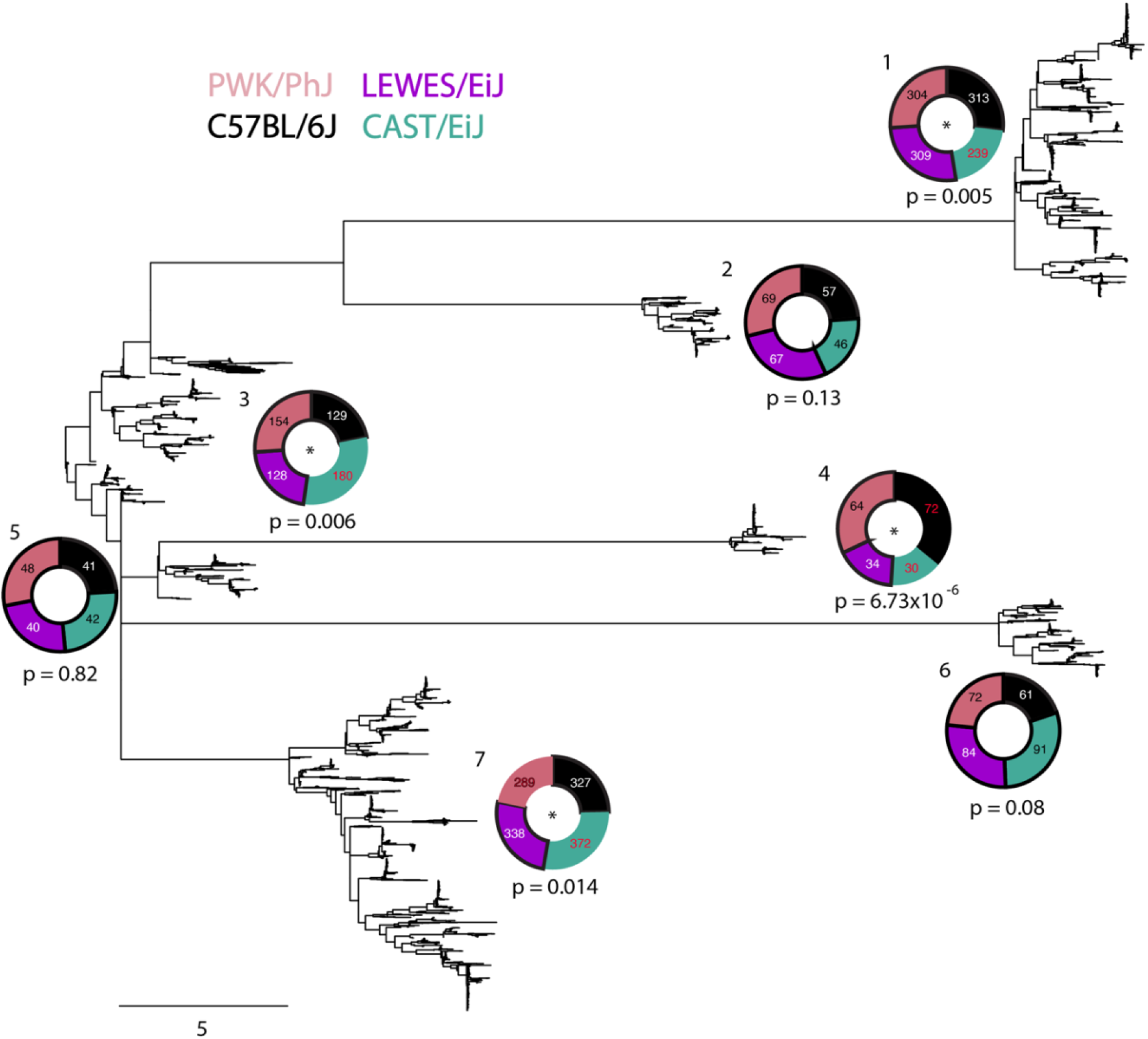
CENP-A associated sequences form distinct subgroups. Neighbor joining tree constructed from each strain’s top 1000 CENP-A associated sequences. Sequences cluster into seven groups. Strain-level contributions to each group are depicted by donut plots. The number of sequences from each strain are given in each wedge of the donut. Groups with skewed strain representation have an asterisk at the center of the donut plot (Groups 1, 3, 4, 7). P-values were calculated using Chi-square tests. For clades with biased strain representation, Bonferroni post hoc tests were used to identify strains driving the significant signal. These strains are represented in the donut plots with red numbers and without a black outline.

### CENP-A associated sequences exhibit phylogenetic similarities across strains

We next sought to holistically assess the similarity among the 1000 most CENP-A enriched sequences in each strain using a computational stylistics analysis. Results from this analysis are summarized in a 2-dimentional PCA plot (Figure 5) that exposes two notable trends.

**Figure 5:**
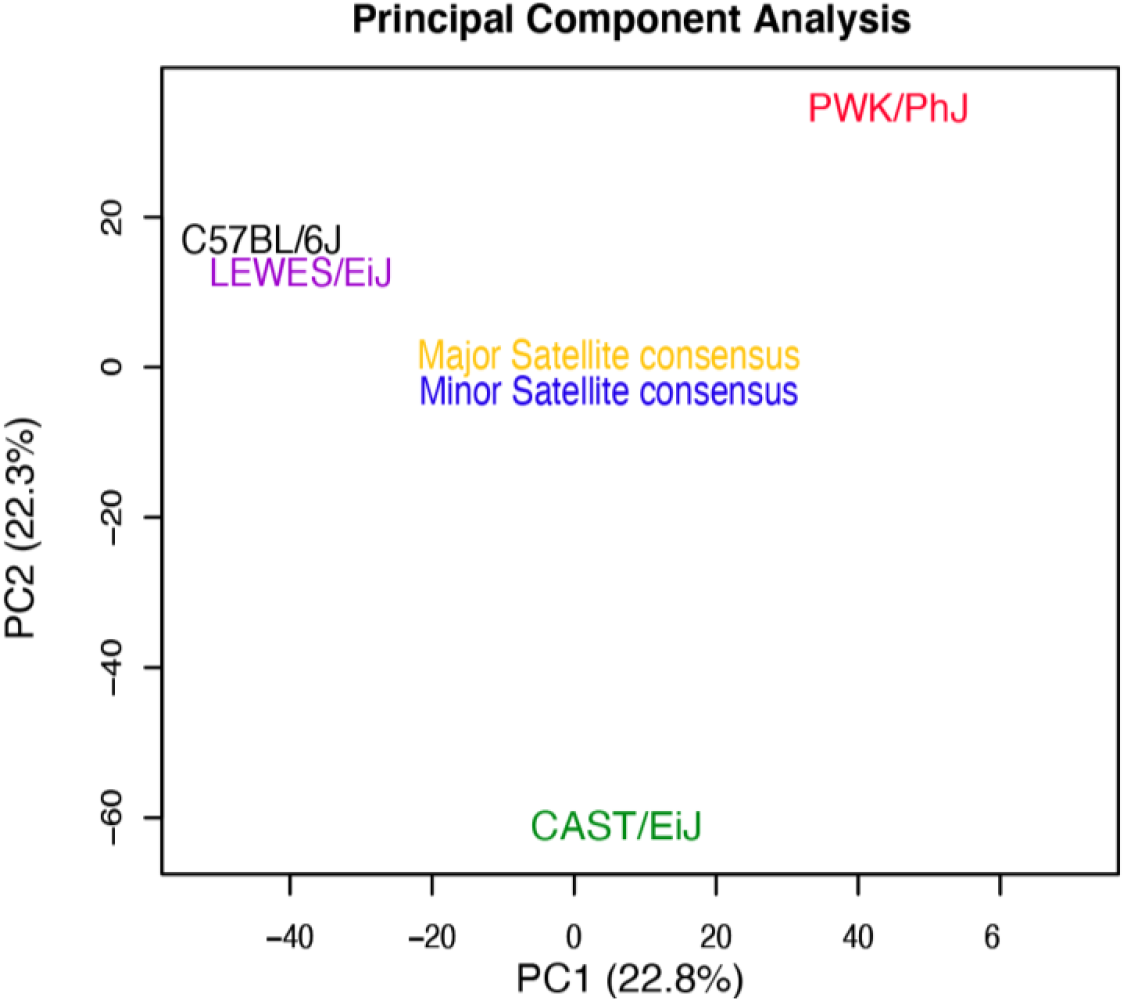
Relationship between CENP-A associated sequences across strains. (A) Principal component analysis derived from computational stylistics analysis of the minor and major satellite consensus sequences and CENP-A associated sequences in each strain.

First, C57BL/6J and LEWES/EiJ, two *M. musculus domesticus* strains, share high similarity in their CENP-A associated sequences. CAST/EiJ and PWK/PhJ have distinct CENP-A associated sequence libraries that partition these strains along unique coordinates in PCA space (Figure 5). While our strain sample size is small, our findings suggest stark subspecies level trends in CENP-A associated centromere satellite sequences. This patterning stands in contrast to the absence of an evolutionary signal in measures of overall centromere sequence and architecture (Arora et al. 2021), suggesting that the CENP-A associated sequence landscape is more slowly evolving than the underlying sequence of the centromere itself.

Second, the minor and major centromere satellite sequences form a focal hub around which the CENP-A associated centromere satellites are organized (Figure 5). This patterning suggests that, while the consensus sequence is an average representation of the CENP-A associated centromere satellites in *M. musculus*, a single sequence cannot comprehensively capture the diversity of centromere satellite sequences present in each strain’s genome.

The top 1000 sequences for each strain were then input into the MEME motif discovery tool (Bailey et al. 2009) to uncover sequence motifs associated with CENP-A in each strain (Supplemental Table S1). Enriched motifs vary between strains and localize to discrete positions along the minor satellite (Supplemental Figure S6). Thus, CENP-A preferentially associates with distinct sequence motifs along the minor satellite repeat in each strain.

### Strain-specific transcription factor motif presence across CENP-A associated sequences

Centromere transcription is essential for differentiation and development and is tightly regulated in both timing and rate (Smurova and De Wulf 2018). Specifically, centromere-derived RNAs are crucial for localizing HJURP, a CENP-A chaperone that guides CENP-A incorporation into nucleosomes, to specify centromere identity. Thus, centromere transcription plays critical roles in early kinetochore assembly, including CENP-A recruitment (Smurova and De Wulf 2018). However, the specific transcription factors (TFs) that act at centromeres to regulate transcriptional activity remain largely unknown (Ferri et al. 2009). We reasoned that strain differences in CENP-A sequence association might stand to offer insight into strain differences in centromere transcription dynamics.

To explore this possibility, we used MEME-ChIP to identify specific TF-binding motifs enriched among strain specific CENP-A associated sequences. We uncovered 105 TF motifs present in at least one strain (Supplemental Figure S7). Thirteen TF motifs were present in all strains (ATF3, BATF, CUX1, CUX2, DLX5, FOSB, HNF1A, HNF1B, HSF2, JUN, JUNB, LHX3, LHX6, PRGR and SOX2), although strains differ in the frequency of these TF motifs. Overall, the frequency and presence of TF motifs is highly variable across the suite of CENP-A associated sequences in each strain.

We next sought to verify whether the 105 observed TF motifs in CENP-A associated sequences lead to binding of a TF with the centromere. We utilized publicly available ChIP-seq data from the NCBI Sequence Read Archive and mapped reads from ChIP and input samples to the centromere satellite consensus. As most ChIP-seq data is from the reference mouse strain C57BL/6J, and since the consensus minor satellite sequence is the most abundant in this strain (Supplemental Figure S2), we reasoned that mapping ChIP-seq reads to the centromere consensus would enable discovery of TF association at the centromere. We classify centromere associated TFs as those with a higher average percentage of mapped reads localizing to the minor satellite in the TF ChIP compared to the input sample.

As expected, we find evidence of both histone H3 (Supplemental Figure S8A) and the histone modification H3K9me3 (Supplemental Figure S8B) at minor and major centromere satellite DNA, respectively, validating our methodology.

We identified two centromere-associated TFs – SMAD3 and HNF1A (Figure 6). Of note, SMAD3 motifs are differentially prevalent in the top 1000 CENP-A associated sequences across our four strains, with the motif altogether absent in CAST/EiJ CENP-A associated sequences (Figure 6A and B). We speculate that the absence of SMAD3 motifs in CAST/EiJ CENP-A associated sequences could lead to strain differences in centromere transcription dynamics in response to SMAD3 signaling.

**Figure 6:**
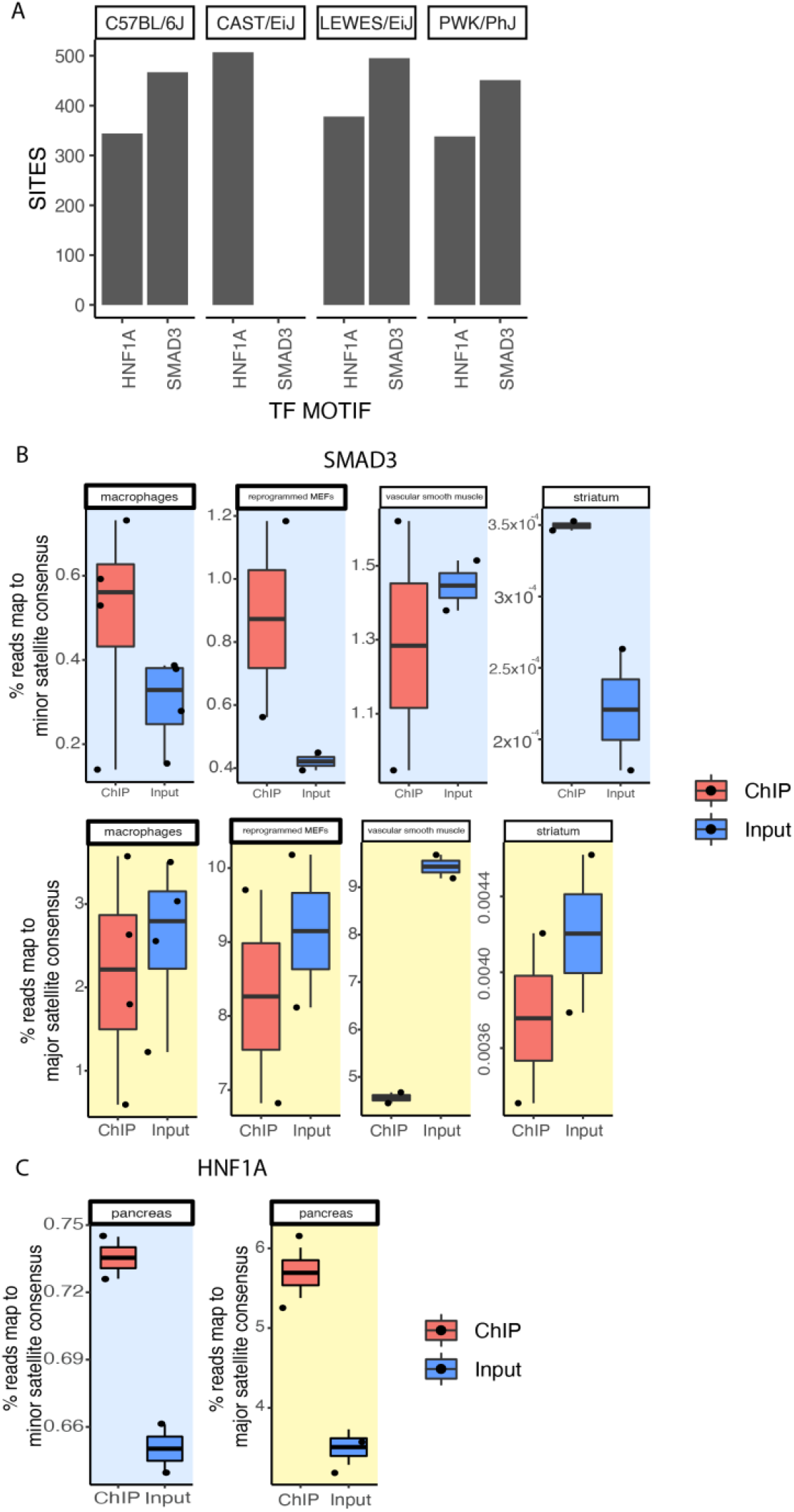
Transcription factor occupancy at centromere satellite DNA. (A) Transcription factor motif frequency in strain-specific CENP-A enriched sequences. (B and C) Association of transcription factors SMAD3 and HNF1A at centromeres from publicly available ChIP-seq data. The y-axis represents the percentage of reads that map to the minor satellite (blue) and major satellite (yellow) consensus sequence in ChIP or input samples (x-axis). Each dot represents the average value of replicates in an experiment.

## DISCUSSION

Rapid centromere sequence evolution imposes complementary selection pressures on centromere-associated kinetochore proteins, leading to dynamic species-level trends in centromere satellite-kinetochore protein co-evolution. Despite this well-established paradigm between species, the fundamental question of how within species genetic diversity at the centromere impacts kinetochore protein association remains unaddressed. Our recent work exposed substantial polymorphism in centromere satellite sequences between inbred house mouse (*Mus musculus*) strains (Arora et al. 2021). In this study, to quantitatively assess the effect of centromere satellite diversity on CENP-A (the centromere-specific histone variant) association, we performed CENP-A ChIP-seq in four diverse inbred mouse strains. Although CENP-A is rapidly and adaptively evolving in some taxa, our four surveyed mouse strains share an identical CENP-A amino acid sequence. Thus, our experimental framework allows us to singularly investigate how centromere DNA diversity impacts the association of CENP-A without introducing confounding effects from protein-level variation.

We uncover pronounced strain differences in the positioning of CENP-A along the minor (functional centromere core) satellite sequence (Figure 2A). CENP-A positioning along the minor satellite trended with minor satellite repeat diversity (Arora et al. 2021), with strains harboring reduced minor satellite repeat diversity exhibiting more similar association profiles (C57BL/6J and PWK/PhJ) than strains with higher minor satellite repeat diversity (CAST/EiJ and LEWES/EiJ). Prior research has suggested that centromere satellites comprised of different repeat structures have variable stability and competence for centromere function (Aldrup-MacDonald et al. 2016). Our findings reinforce the possibility that centromere repeat diversity influences CENP-A spacing and density, properties that could, in turn, have consequences for the architecture and stability of the kinetochore complex (Walstein et al. 2021).

Canonically, CENP-A is thought to associate with the minor satellite array located in the functional centromere core. The minor satellite array is therefore predicted to be the primary determinant of centromere size in house mice. In contrast to this widely held belief, we find significant enrichment of CENP-A in the pericentromeric major satellite in inbred strains CAST/EiJ and LEWES/EiJ (Figure 1B). These observations reveal a possible role of the major satellite array in the regulation of centromere size and imply that functional centromere size may not always be strictly proportional to the length of the minor satellite array. The major satellite array is known to recruit microtubule destabilizers, which enable stronger centromeres to reorient towards to oocyte during female meiosis (Kumon et al. 2021). Evidence of CENP-A association with the major satellite brings into question its influence on microtubule destabilizer recruitment by the major satellite and the propensity for centromere drive. Given the established relationship between centromere size and centromere drive potential, our results suggest that strain differences in CENP-A association with pericentromeric chromatin could lead to genotype-dependent differences in chromosome transmission dynamics.

Although we consider only one replicate of H3K4me3 per strain, there is a concomitant association of H3K4me3 with the major satellite in strains that show evidence for CENP-A association with the major satellite. This finding raises further questions about the relationship between epigenetic modifications at canonical histones and CENP-A recruitment at pericentromeric chromatin, as well as the potential for strain differences in the epigenetic landscape of centromeres.

CENP-A association was previously shown to center on the 17-bp CENP-B motif in the C57BL/6J and ZALENDE/EiJ inbred mouse strains (Iwata-Otsubo et al. 2017). We replicate this earlier finding for C57BL/6J and further show that CENP-A binding localizes to the CENP-B box in PWK/PhJ (Figure 2A). However, two other profiled strains in our study – LEWES/EiJ and CAST/EiJ – show no evidence of preferential association at this locus. CENP-B is a constitutively expressed and conserved DNA binding protein but, paradoxically, is not required for chromosome segregation (Hudson et al. 1998). Despite its non-essentiality, CENP-B has multiple centromere-associated functions, including the recruitment and stabilization of CENP-A, regulation of centromeric heterochromatin, and de novo centromere formation (Gamba and Fachinetti 2020). We show that these strain differences in CENP-A localization to the CENP-B box are not due to variation in the relative abundance of CENP-B boxes across strains (Figure 2B), suggesting that they reflect underlying strain differences in CENP-A affinity for the CENP-B motif. Whether this differential affinity is, in turn, mediated by strain differences in CENP-B binding at the CENP-B motif remains unknown.

We previously observed that shared evolutionary history is a poor predictor of minor satellite copy number and sequence heterogeneity amongst inbred mouse strains, suggesting that minor satellite arrays evolve sufficiently rapidly to outstrip signals of recent shared descent amongst inbred mouse strains (Arora et al. 2021). Although we are underpowered to detect clear phylogenetic signals in this study, we observe that more closely related strains exhibit more similar CENP-A associated sequence profiles (Figure 5). This observation suggests that while the minor satellite array is evolving rapidly to outstrips signals of phylogenetic relatedness, CENP-A association with the minor satellite array might follow a phylogenetic pattern.

Our findings establish pronounced differences in CENP-A sequence localization among inbred mouse strains. CENP-A localization and integration into centromeric histones is dependent on RNA Polymerase II-mediated centromere transcription. Additionally, centromere transcription plays key roles in cellular proliferation and differentiation (Smurova and De Wulf 2018). We report evidence for strain differences in TF motif presence at CENP-A associated sequences, suggesting strain variation in centromere transcriptional response. As a notable example, we find SMAD3 binding sites in C57BL/6J, LEWES/EiJ, and PWK/PhJ but not CAST/EiJ CENP-A associated sequences. Through re-analysis of published SMAD3 ChIP-seq data from C57BL/6J, we show that SMAD3 binding enrichment is limited to the minor satellite (to the exclusion of the major satellite) across multiple cell types. SMAD3 is a transcription factor that regulates cellular proliferation in response to extracellular cues (Roberts et al. 2003), particularly in the context of wound healing and cancer. The absence of SMAD3 motifs at CAST/EiJ CENP-A associated sequences could contribute to strain differences in cellular proliferation with consequences for regeneration or the progression of cancer. Overall, these centromere-mediated transcriptional differences between strains could provide a mechanism for individual differences in genome stability in response to external cues.

In addition to uncovering novel centromere biology, our work also pioneers the application of powerful reference genome independent strategies for probing the epigenetic and TF landscape of repetitive sequences refractory to analysis with conventional methods. Broader application of these methods to other ChIP-seq experiments will enhance understanding of the chromatin environment of centromeres, as well as other regions of the genome that are currently absent from the reference assembly. Such analyses will yield a more comprehensive picture of chromatin regulation across the genome and offer new functional insights into the most intractable genomic regions.

Understanding how centromere variation influences genome stability and chromosome transmission dynamics remains a key challenge for the field of centromere biology. Here, we hypothesized that centromere variation can exert functional effects via altering association with kinetochore proteins. Using an innovative and customized *k*-mer based, reference-blind bioinformatics strategy, we show that centromere diversity in four diverse inbred mouse strains is associated with dramatic differences in the association of one crucial kinetochore protein, CENP-A. We further show that these strain differences in CENP-A association are linked to variation in TF motif presence and likely differences in centromere transcription dynamics. Importantly, the discovery of individual level differences in the centromere association landscape of an essential kinetochore protein presents a novel mechanism for variation in genome stability and chromosome segregation dynamics, phenomena that may underpin the genetic etiology cancer and infertility.

## MATERIALS AND METHODS

### Mice

C57BL/6J (stock no 000664), LEWES/EiJ (stock no 002798), CAST/EiJ (stock no 000928), and PWK/PhJ (stock no 003715) mice were obtained from The Jackson Laboratory. Mice were housed in a low barrier room and provided food and water *ad libitum*. Mice were euthanized by CO_2_ asphyxiation or cervical dislocation in accordance with recommendations from the American Veterinary Medical Association. All animal experiments were approved by the Institutional Animal Care and Use Committee and were consistent with the National Institute of Health guidelines.

### Chromatin extraction

Chromatin was extracted from two or three flash-frozen mouse livers per CENP-A ChIP using a previously described protocol (Iwata-Otsubo et al. 2017). Livers were homogenized in 4 mL ice-cold Buffer I (0.32 M Sucrose, 60 mM KCl, 15 mM NaCl, 15 mM Tris-Cl pH 7.5, 5 mM MgCl_2_, 0.1 mM EGTA, 0.5 mM DTT, 0.1 mM PMSF, 50 uL PIC (Sigma P8340) per g of tissue). The homogenate was filtered through a 100 μm cell strainer (Fisher) and centrifuged at 6000g for 10 min at 4°C. The pellet was resuspended in the same volume of Buffer I and an equivalent volume of ice-cold Buffer I supplemented with 0.2% NP-40 alternative (Sigma) was added. Samples were incubated on ice for 10 min to release nuclei. 4mL of nuclei were gently laid upon 16 mL of ice-cold Buffer III (1.2 M Sucrose, 60 mM KCl, 15 mM NaCl, 15 mM Tris-Cl pH 7.5, 5 mM MgCl_2_, 0.1 mM EGTA, 0.5 mM DTT, 0.1 mM PMSF, 50 uL PIC (Sigma P8340) per g of tissue) in a 50 mL conical tube. Samples were centrifuged at 10,000g for 20 min at 4°C. The supernatant was discarded, and the pellet was resuspended in 26 uL MNase buffer (50 mM Tris, 1 mM CaCl_2_, 4 mM MgCl_2_, 4% NP-40) supplemented with 1 mM PMSF and 1:100 dilution of PIC (Sigma P8340) per 10×10^6^ cells. Cell number was calculated invoking the assumption that 1 g of mouse liver tissue is equivalent to 125 million cells (Sohlenius-Sternbeck 2006). MNase (100U/uL Thermo PI88216) was added at a concentration of 1uL/33.3×10^6^ cells and samples were incubated at 37°C for 12 minutes. MNase digestion was stopped by adding 0.5M EDTA to a final concentration of 10 mM. Samples were incubated on ice for 5 min and then spun at 15000 rpm for 10 min. The supernatant was transferred to a new tube and spun again at 15000 rpm for 10 min. Finally, the supernatant was transferred to a new tube and either stored at −20°C or carried forward into chromatin immunoprecipitation.

### Chromatin immunoprecipitation (ChIP)

ChIP was performed using an antibody against CENP-A (D601AP developed by Beth Sullivan, Duke University) and H3K4me3 (Millipore Cat. # 07-473) (Maloney et al. 2012; Iwata-Otsubo et al. 2017). 25 uL Protein G dynabeads (Invitrogen Cat. #10003D) were used per ChIP reaction. Beads were washed two times with RBD (RIPA buffer (Sigma Cat. # R0278), 50 mg/mL Bovine Serum Albumin (BSA), and 0.5 mg/mL Herring Sperm DNA). Beads were resuspended in RBD, combined with antibody, and then allowed to conjugate at room temperature for 20 min with gentle rotation. Again, beads were washed two times in RBD and resuspended in RBD supplemented with 1 mM PMSF and 1:100 dilution of PIC (Sigma P8340). Chromatin was added to tubes and allowed to incubate at 4°C with rotation overnight. A 10x chromatin volume was used for CENP-A ChIP and a 2x chromatin volume was used for H3K4me3 and IgG ChIP compared to input chromatin. Beads were then washed three times with RIPA buffer and 50 mg/mL BSA. Beads were subsequently washed two times with TE pH 8.0. Beads were washed once more with TE pH 8.0 and transferred to a new tube. Elution Buffer (1% SDS, 20 mM Tris-HCl pH 8.0, 200 mM NaCl, 5 mM EDTA) supplemented with Proteinase K (New England Biolabs Cat. # P8107S) was added to the beads for both ChIP and input chromatin. Samples were incubated at 68°C overnight with vigorous shaking in a thermomixer. Chromatin from ChIP samples was recovered from beads using a magnet and placed in a clean tube. Samples were processed using the GeneJET PCR purification kit and eluted in 50 uL of 10 mM Tris-HCl pH 8.0. DNA concentrations were measured using Qubit dsDNA High Sensitivity kit (Thermo Cat. # Q32854) according to the manufacturer’s instructions.

### qPCR validation of ChIP efficacy

Quantitative PCR (qPCR) was performed on a ViiA 7 real-time PCR system (Thermo Fisher). PCR was carried out for 40 cycles followed by a melt curve analysis. qPCR was performed on all replicate samples from CENP-A, H3K4me3, and IgG ChIP and input experiments. We measured the relative cycle number (determined by automated threshold analysis by the machine) for ChIP compared to input samples for each ChIP reaction and primer pair. Reactions were run with PowerUp SYBR Green 2x master mix and the following primers - ActB Promoter F primer: 5’-GCCATAAAAGGCAACTTTCG-3’, ActB Promoter R primer – 5’-TTTCAAAAGGAGGGGAGAGG-3’, Minor Satellite qPCR F: 5’-CAT GGAAAATGATAAAAACC-3’, Minor Satellite qPCR R: 5’ -CATCT AATATGTTCTACAGTGTGG-3’.

### Library preparation and sequencing

ChIP libraries were constructed using the KAPA HyperPrep Kit (Roche Sequencing and Life Science) according to the manufacturer’s protocols. Briefly, the protocol entails ligating Illumina specific barcoded adapters, size selection, and PCR amplification. The quality and concentration of the libraries were assessed using the High Sensitivity D5000 ScreenTape (Agilent Technologies) and KAPA Library Quantification Kit (Roche Sequencing and Life Science), respectively, according to the manufacturers’ instructions. Libraries were sequenced using 75 bp single-end reads on an Illumina NextSeq 500 using the High Output Reagent Kit v2.5.

### Centromere consensus read mapping analysis

We used fastp (version 0.23.1) for preprocessing of fastq files to filter low quality reads and trim adapter sequences (Chen et al. 2018). Reads were then mapped to the major and minor satellite centromere consensus sequences using bwa (version 0.7.9) (Li and Durbin 2010). The percentage of reads that mapped to each consensus sequence was calculated from output of the idxstats command in samtools (version 1.8) and the total number of reads in the fastq file (Li et al. 2009). Enrichment at centromeres was quantified relative to input, normalizing by the number of sequenced reads. To visualize the relationship between enriched mapped sequences across strains, we constructed a neighbor joining tree with R package ape (version 5.6-2) in R (version 4.0.5).

### *k*-mer tables and strain-specific enriched *k*-mers

Fastp-filtered reads were processed through clumpify (https://sourceforge.net/projects/bbmap) to remove optical duplicates. This step ensured that *k*-mer counts were not skewed by PCR-related artifacts. Each sequenced read in a sample’s fastq file was then decomposed into its constituent nucleotide words of length *k*, or *k*-mers, using a custom Python script (KmerComposition.py). We set *k*=31 to balance the competing demands of computational resource usage and sequence specificity.

*k*-mer frequencies were normalized to the number of reads. We then calculated an “enrichment score” for each *k*-mers as the ratio of the normalized *k*-mer frequency in CENP-A ChIP compared to input samples. For each strain, *k*-mers were then ranked by the enrichment score to identify the top 0.1% most enriched *k*-mers.

To identify *k*-mers with upto 4 mismatches with the consensus minor satellite sequence, we mapped whole genome *31*-mers to the minor satellite consensus sequence by invoking the *aln* command in bwa version 0.7.9 (Arora et al. 2021). We looked for overlap between the whole genome *31*-mers with upto 4 mismatches to the minor satellite consensus sequence and each strains CENP-A enriched *31*-mers.

### Scoring CENP-A ChIP-seq reads for enrichment of *k*-mers and abundance

To prioritize a list of highly enriched CENP-A associated sequences in each strain, we assigned two numerical scores to each CENP-A ChIP read. First, we tallied the number of 0.1% CENP-A ChIP/input enriched *k*-mers observed in a given sequencing read from that strain (‘ read score’). We then counted the frequency of each 75-bp sequencing read in each library, normalizing to the total number of sequencing reads (‘read count’). We then jointly ranked sequences by their read score and read count and focused on the top 1000 ranking reads.

### Analyzing strain-specific sequence groups

To visualize the relationship between sets of CENP-A enriched sequences across strains, we constructed a neighbor joining tree using MEGA11 and analyzed subclusters of sequences with R package ape (version 5.6-2) in R (version 4.0.5).

To compare different sets of sequences across strains, we performed a computer-assisted text analysis using the package stylo (version 0.7.4) in R (version 4.0.5). Relatedness among groups of words (in our case strain-specific top 1000 ranked 75-bp reads) was assessed via a principal component analysis. Principal component analysis was performed using the 3872 most frequent words culled at 0%. 1000 bootstrap cluster analyses with randomly sampled sets of 1000 sequences were also performed and confirmed *P* < 0.001 significance of top 1000 ranked read results. Relatedness among strain-specific top ranked 75-bp reads and the minor and major satellite consensus sequences were assessed via a principal component analysis performed using the 5329 most frequent words culled at 0%.

### MEME motif enrichment analysis

We used the MEME and MEME-ChIP tools in the MEME suite (v5.4.1) to identify specific motifs enriched in CENP-A associated sequences.

MEME is a motif discovery algorithm that employs a probabilistic model. Each discovered motif is assigned an E-value corresponding to the expected number of motifs with the given log likelihood ratio (or higher), and with the same width and site count, that one would find in a similarly sized set of random sequences (sequences where each position is independent, and letters are chosen according to the background letter frequencies). MEME was run separately on the top 1000 CENP-A associated sequences for each strains by using the following command “meme STRAIN_top1000.fa -dna -oc . -nostatus -time 14400 -mod zoops -nmotifs 3 -minw 6 -maxw 50 -objfun classic -revcomp -markov_order 0”.

MEME-ChIP performs comprehensive motif analysis on a set of sequences, assuming that motifs are centrally located within the input sequence. MEME-ChIP was run on the top 1000 most CENP-A enriched sequences in each strains with the following command “meme-chip -oc . -time 240 -ccut 100 - fdesc description -dna -order 2 -minw 6 -maxw 15 -db db/MOUSE/HOCOMOCOv11_core_MOUSE_mono_meme_format.meme -meme-mod zoops -meme-nmotifs 3 -meme-searchsize 100000 -streme-pvt 0.05 -streme-totallength 4000000 -centrimo-score 5.0 - centrimo-ethresh 10.0 STRAIN_top1000.fa”

### Transcription factor association analysis

ChIP-seq data from 105 identified transcription factors (TFs) were obtained in fastq format from the NCBI Sequence Read Archive (SRA). Reads were then mapped to the major and minor satellite centromere consensus sequences using bwa version 0.7.9 (Li and Durbin 2010). As above, the percentage of reads that mapped to each consensus sequence was calculated from output of the idxstats command in samtools (version 1.8) and the total number of reads in the fastq file (Li et al. 2009). We restricted our focus to TFs that (1) had both ChIP and input sequencing samples to enable comparisons of average relative TF enrichment across independent studies and (2) featured ChIP-seq data from more than one study in a particular cell type. Sample information was annotated using the metadata from SRA for each experiment. Data used in this analysis is from BioProject and accesssion numbers in Supplemental Table S2.

## DATA ACCESS

The CENP-A ChIP-seq data generated in this study have been submitted to the NCBI BioProject database (https://www.ncbi.nlm.nih.gov/bioproject/) under accession number PRJNA838487. The scripts and processed files used to analyze data in this study are publicly available through github (https://github.com/umaarora/CENPA-ChIP).

## COMPETING INTEREST STATEMENT

The authors declare no competing interests.

## ACKNOWLEDGEMENTS

We gratefully acknowledge the contribution of the Genome Technologies Service at The Jackson Laboratory for expert assistance with the work described in this publication. We thank Dr. Christopher Baker and Catrina Spruce for protocols to perform the chromatin immunoprecipitation experiment. We also thank members of the Dumont lab, and Drs. Mary Ann Handel and Christopher Baker for comments on the manuscript.

This work was funded by a Maximizing Investigators’ Research Award from the National Institute of General Medical Sciences to BLD (R35 GM133415). UA is supported by a Ruth L. Kirschtein Predoctoral Individual Fellowship from the National Cancer Institute (F31CA268727). CENP-A antibody production and affinity purification was supported through NIH grants R01 GM124041 and R01 GM129263 awarded to BAS. The content of this manuscript is the sole responsibility of the authors and does not necessarily represent the official views of the National Institutes of Health.

## AUTHOR CONTRIBUTIONS

UA performed the experiment and analyses. UA and BLD designed the analysis, interpreted the bioinformatic results, and prepared the manuscript. BAS contributed key reagents. All authors read and approved the final manuscript.

---

**Supplemental Figure S1:**
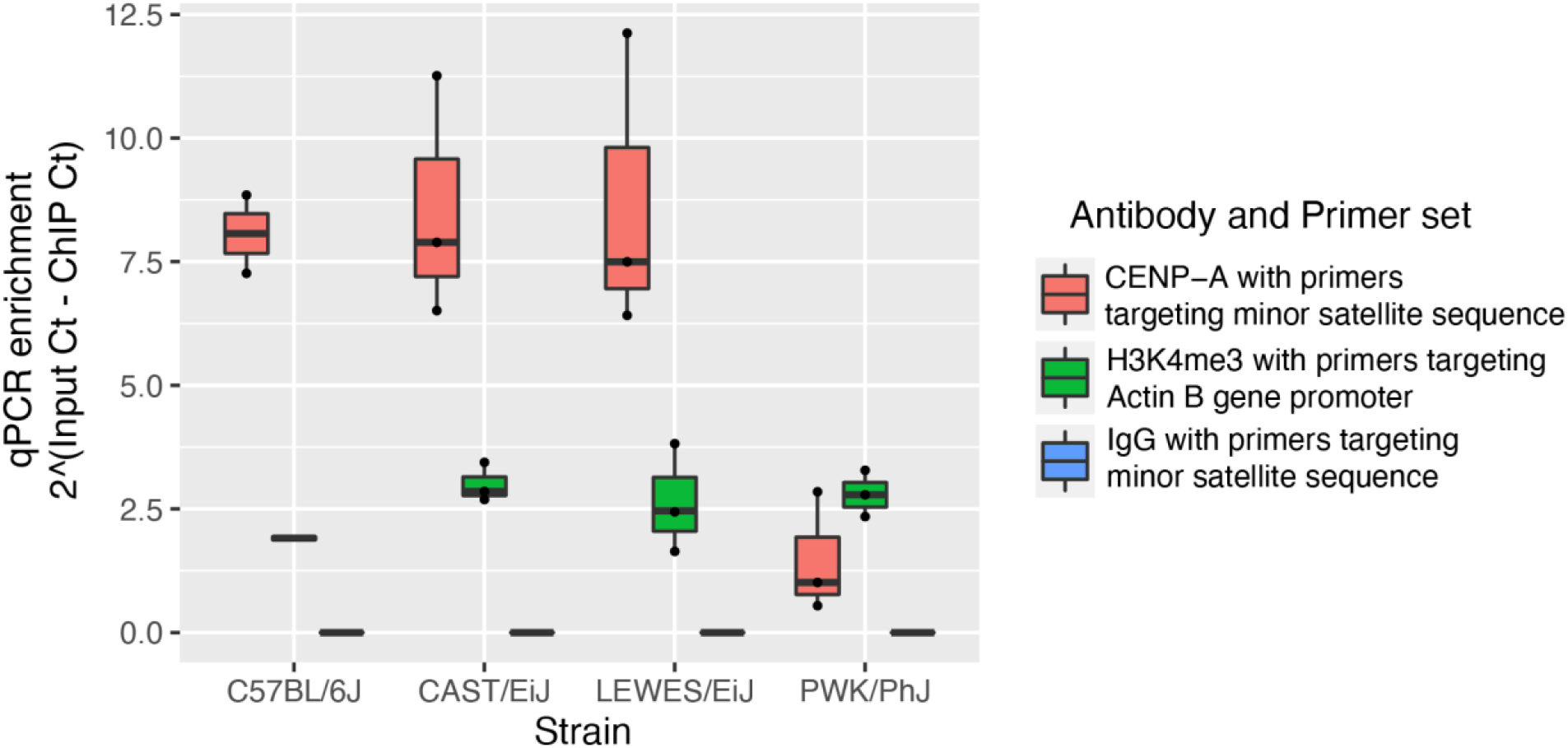
qPCR validation of ChIP DNA before sequencing. Quantitative PCR performed on samples from CENP-A, H3K4me3, and IgG ChIP experiments to validate efficacy. A primer set targeting the minor satellite consensus sequence was used to amplify CENP-A and IgG ChIP DNA. Primers in the Actin B gene promoter were used to amplify H3K4me3 ChIP DNA. Enrichment was calculated using the difference in cycle number between ChIP and input samples. Three replicates were performed for each sample and each strain. For each boxplot, the horizontal line represents the mean, the top and bottom of the box represent the upper and lower quartile respectively, and the vertical line represents the range of values.

**Supplemental Figure S2:**
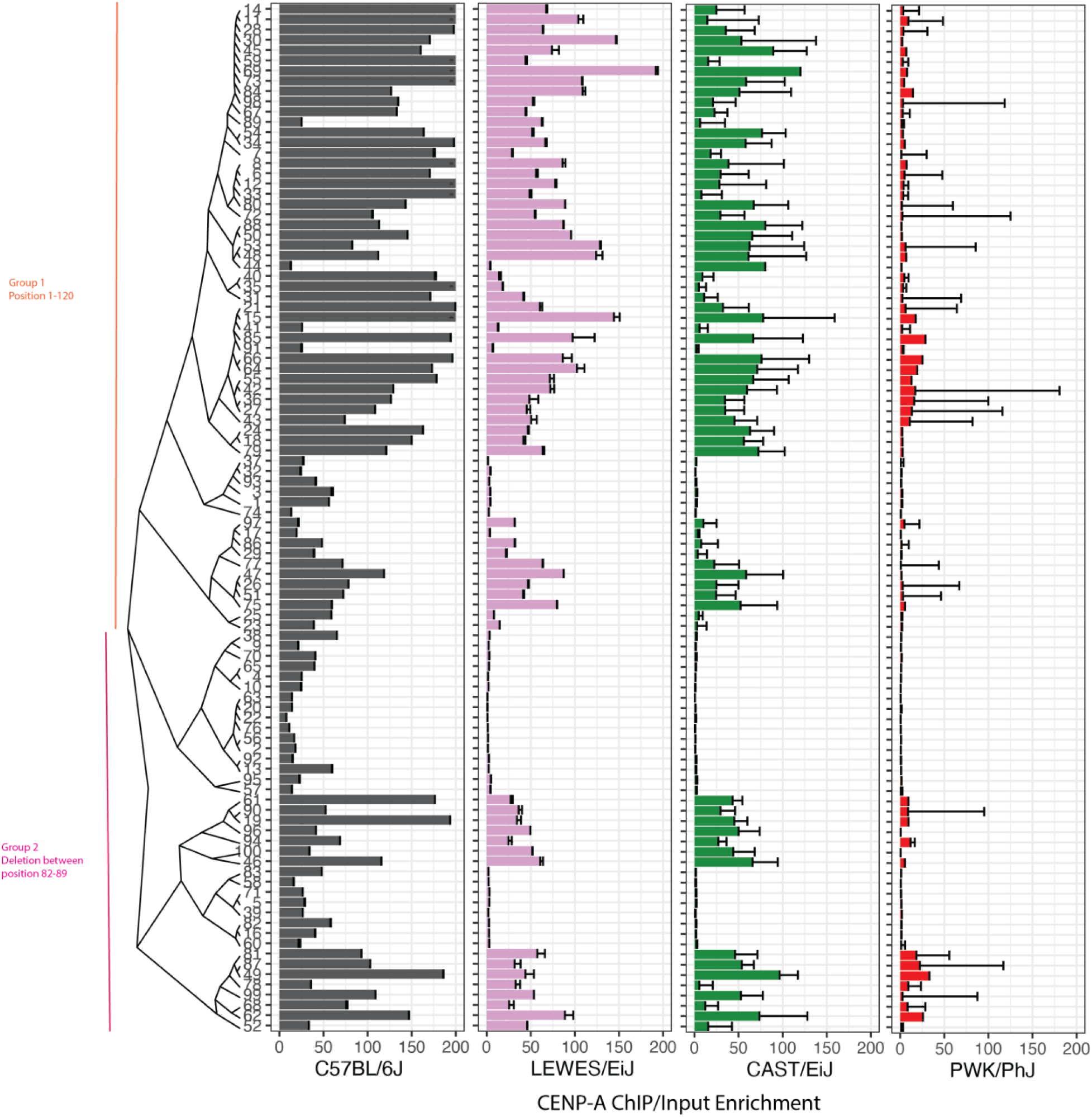
The most differentially abundant CENP-A associated sequences across strains span the minor satellite consensus sequence. Neighbor joining tree depicts the relationship among sequenced reads that map to the minor satellite. These sequences can be broadly categorized into two groups: one matching the consensus sequence and a second carrying a deletion at positions 82-89 relative to the consensus sequence. Each bar chart represents the CENP-A ChIP enrichment of reads for an inbred strain.

**Supplemental Figure S3:**
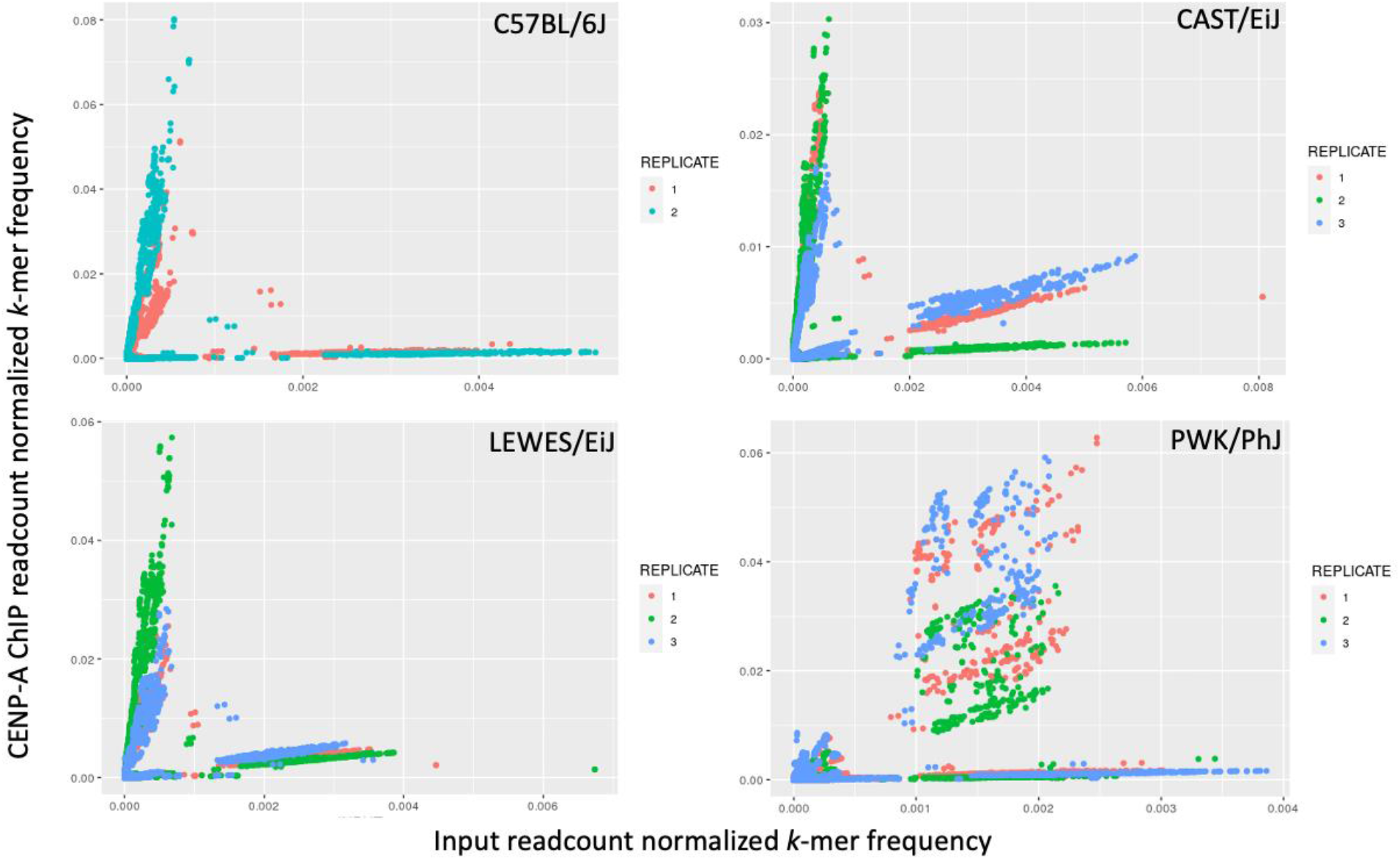
*31*-mer frequency in CENP-A ChIP and input across replicate samples for each strain. Scatter plots representing the normalized abundance of *k*-mers from CENP-A ChIP (y-axis) and input (x-axis) sequencing data for each strain. Replicates are distinguished by color.

**Supplemental Figure S4:**
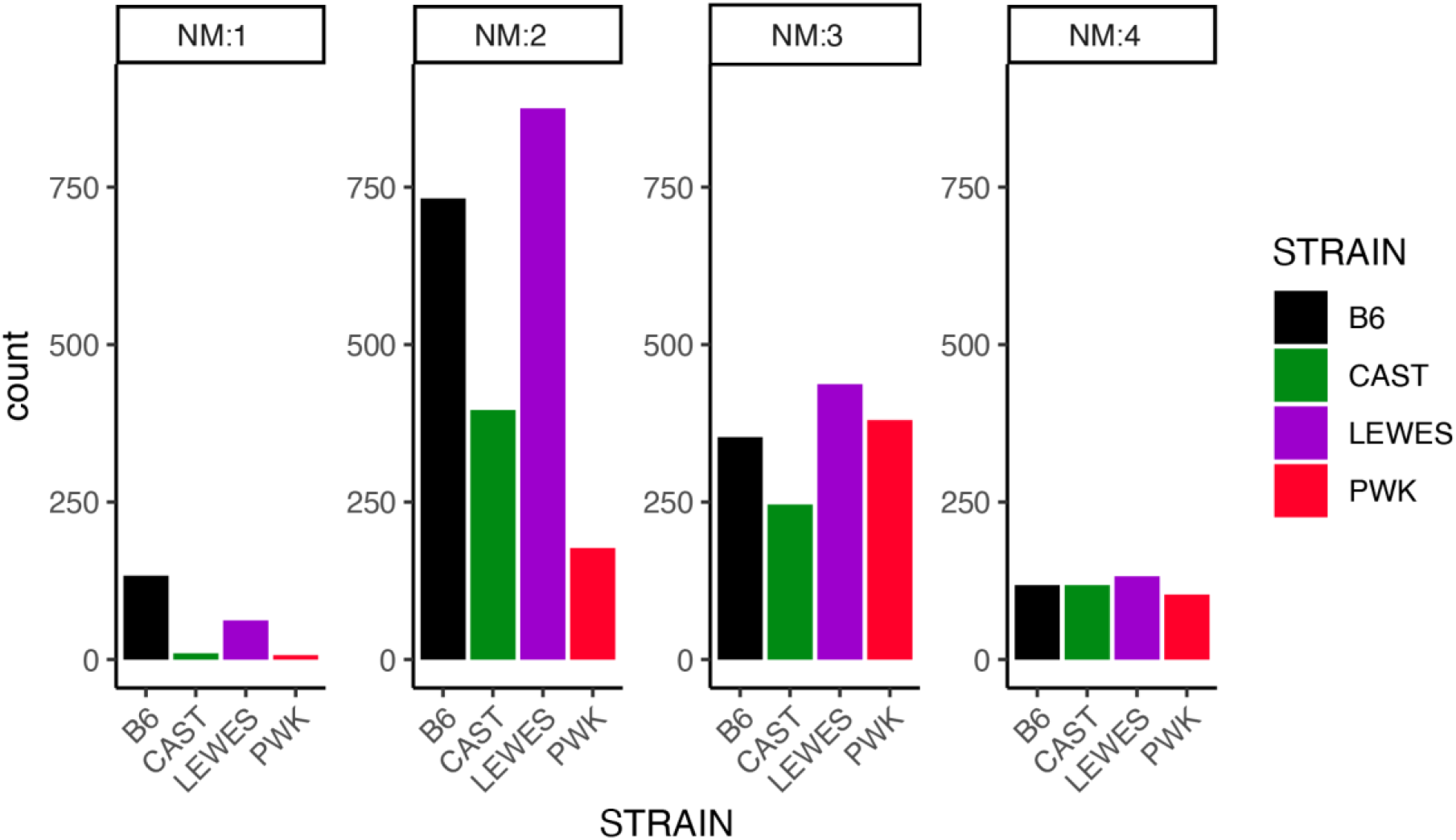
CENP-A enriched *k*-mers carry one or more mismatches from the minor satellite consensus sequence. Bar plots representing the number of CENP-A enriched *k*-mers from each strain with a given number of mismatches (NM) relative to the minor satellite consensus sequence.

**Supplemental Figure S5:**
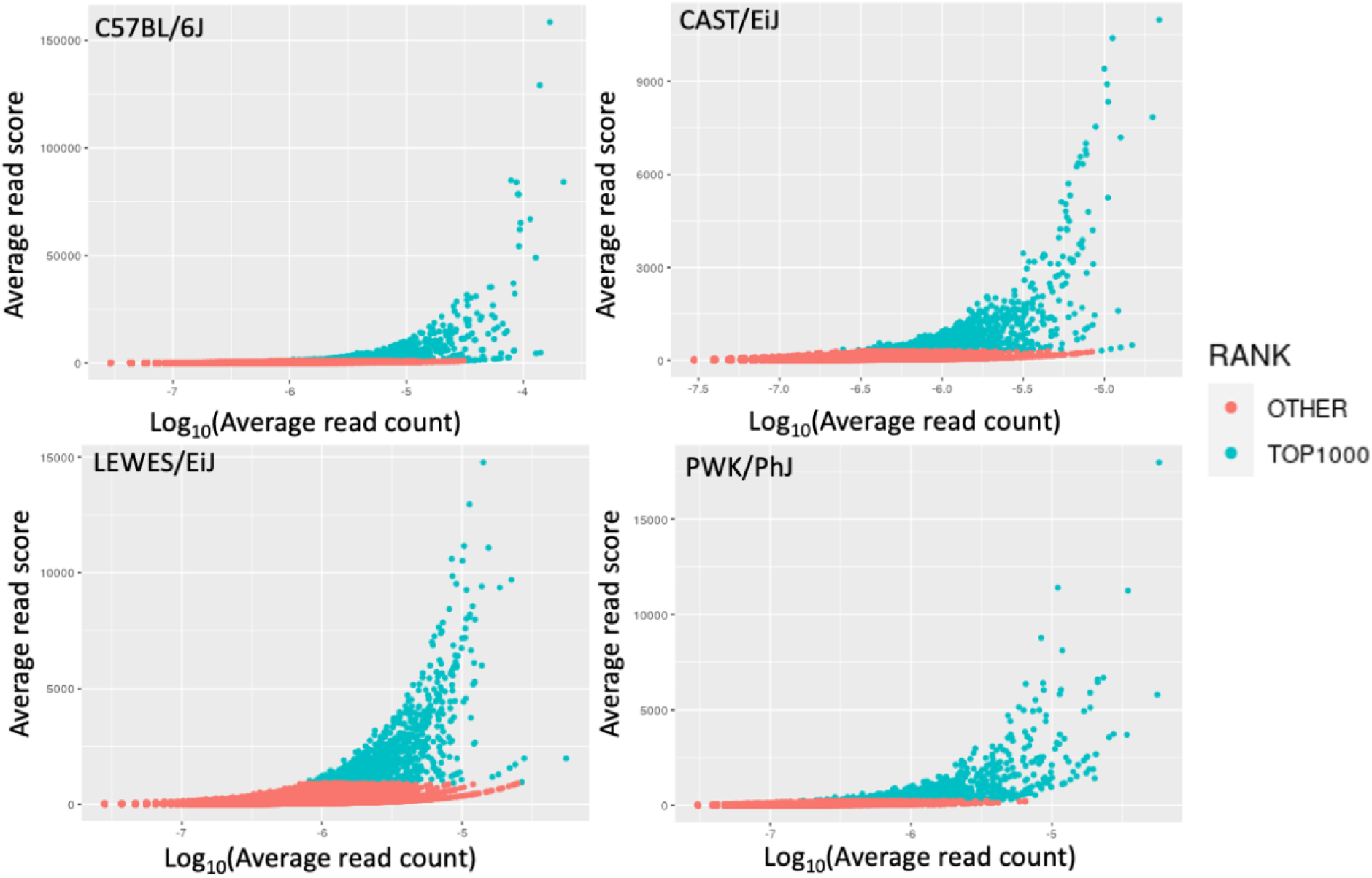
Identification of the top 1000 CENP-A associated sequences for each strain. CENP-A ChIP sequencing reads scored by the frequency of strain-specific *31*-mers (y-axis) and abundance of that read (x-axis) in each strain. Color distinguishes top 1000 scored reads from the remaining reads.

**Supplemental Figure S6:**
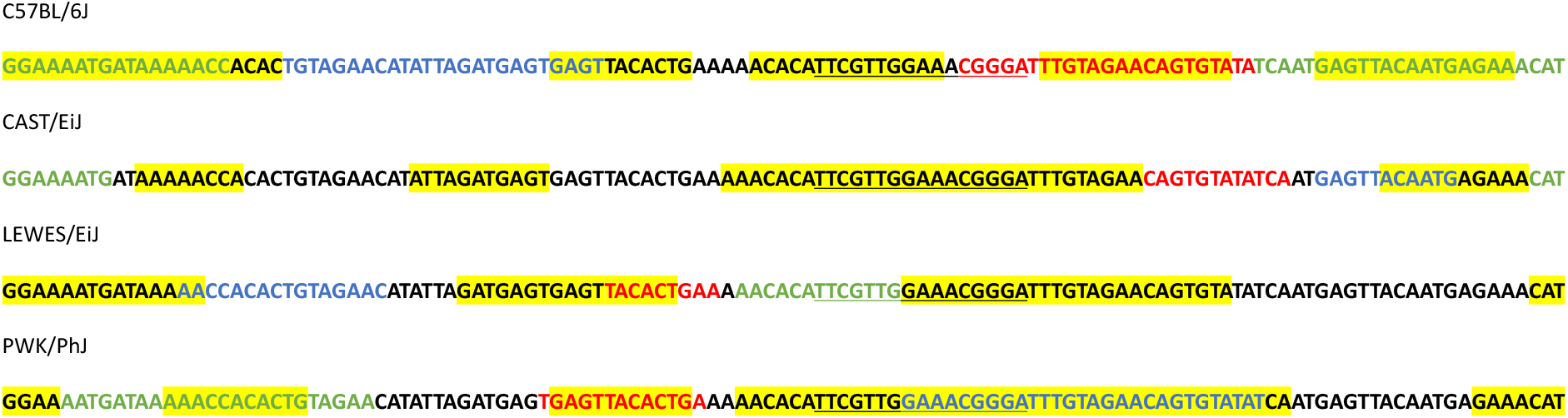
10 most significant motifs from each strain’s top 1000 CENP-A associated sequences. Each line of sequence represents the 120 bp consensus minor satellite repeat unit. The top three CENP-A associated motifs discovered by MEME are highlighted in red, blue, or green. The remaining seven motifs are highlighted in yellow. The 17-bp CENP-B box motif is underlined.

**Supplemental Figure S7:**
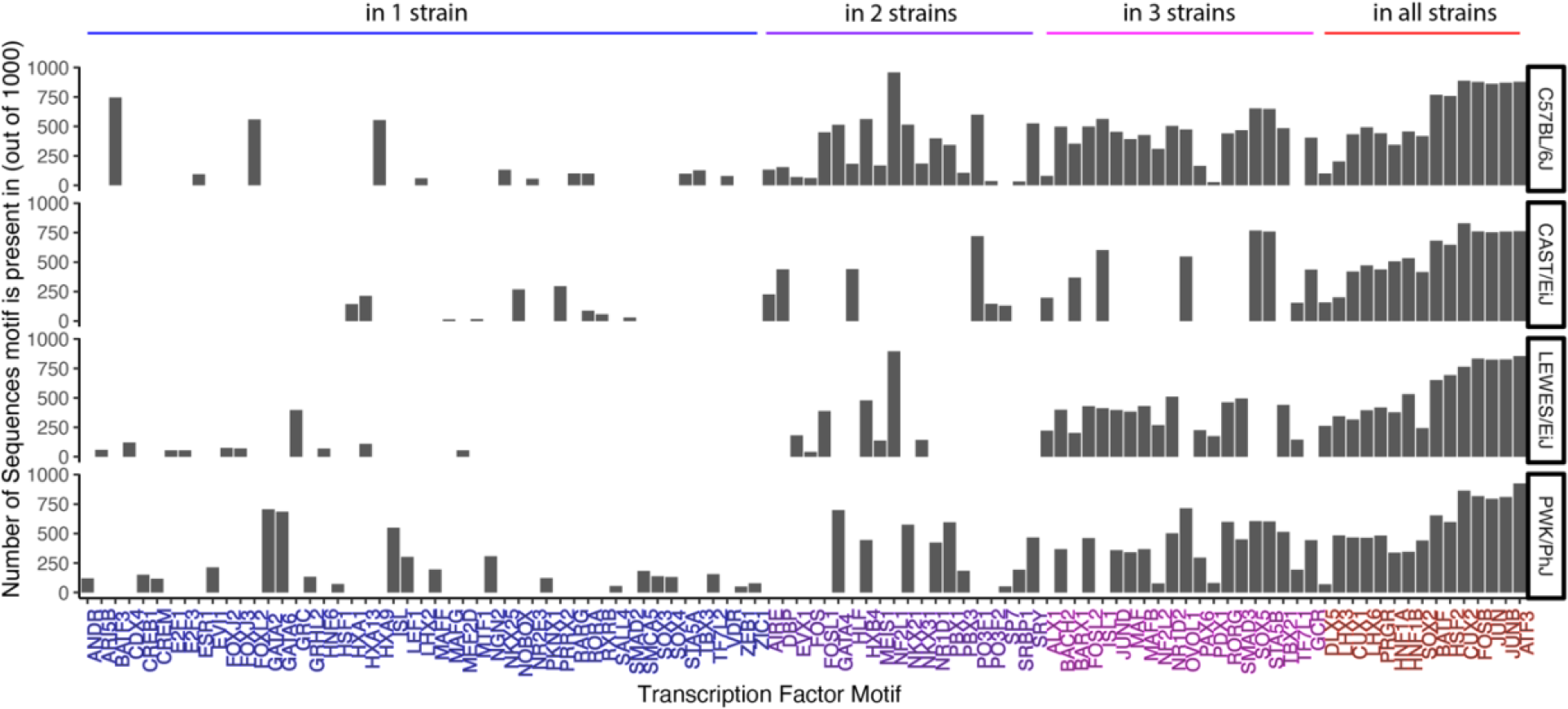
Transcription factor motifs discovered in the top 1000 strain-specific CENP-A enriched sequences. Transcription factor motifs (x-axis) identified in the top 1000 CENP-A associated sequences (y-axis) in each strain (rows). Transcription factors ordered by the number of strains in which the motif is observed.

**Supplemental Figure S8:**
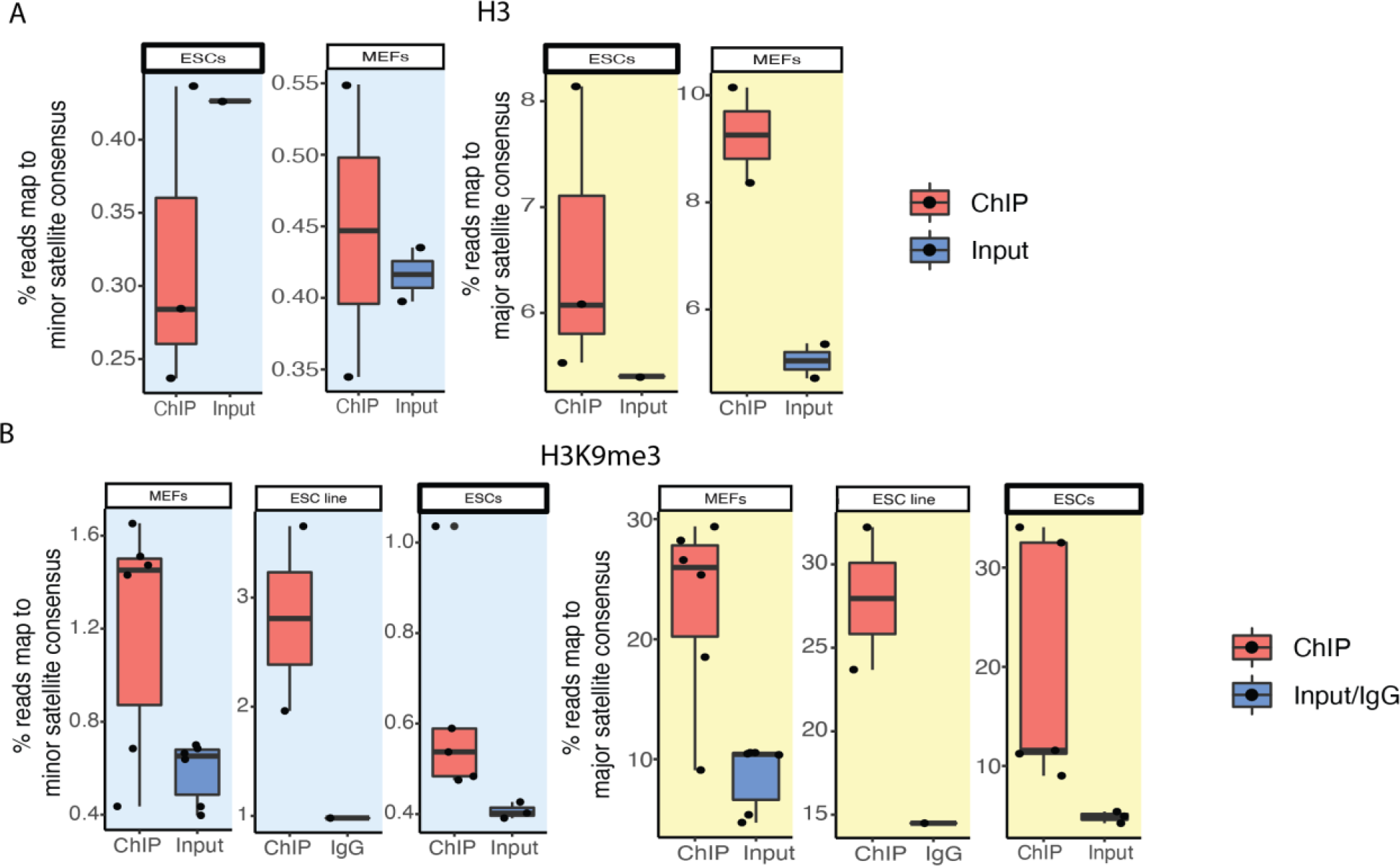
Histone protein occupancy at centromere satellite DNA as a validation for the method to identify transcription factor association at centromere DNA. Association of histones (A) H3 and (B) H3K9me3 at centromeres from publicly available ChIP-seq data. The y-axis represents the percentage of reads that map to the minor satellite (blue) and major satellite (yellow) consensus sequence in ChIP or input samples (x-axis). Each dot represents the average value of replicates in an experiment.

## Notes

### Competing Interest Statement

The authors have declared no competing interest.

